# Discovery of deep-sea coral symbionts from a novel family of marine bacteria, Oceanoplasmataceae, with severely reduced genomes

**DOI:** 10.1101/2022.10.07.511369

**Authors:** Samuel A. Vohsen, Harald R. Gruber-Vodicka, Nicole Dubilier, Charles R. Fisher, Iliana B. Baums

## Abstract

Microbes perform critical functions in corals yet most knowledge is derived from the photic zone. Here, we discovered two mollicutes that dominate the microbiome of the deep-sea octocoral, *Callogorgia delta,* and reside in the mesoglea. These symbionts were abundant across the host’s range, absent in the water, and rare in sediments. The symbionts lack all known fermentative capabilities including glycolysis and can only generate energy from arginine provided by the coral host. Their genomes feature extensive mechanisms to interact with foreign DNA which may be indicative of their role in symbiosis. We erect the novel family Oceanoplasmataceae which includes these symbionts and others associated with four marine invertebrate phyla. Its exceptionally broad host range suggests that the diversity of this enigmatic family remains largely undiscovered. Oceanoplasmataceae genomes are the most highly reduced among mollicutes providing new insight into their reductive evolution and the roles of coral symbionts.

## Introduction

Corals are foundation species that support diverse animal communities from shallow waters to the deep sea. Corals also associate with a diversity of microbes^1^ including the well-studied algal symbionts of the family Symbiodiniaceae and other microbial taxa that perform a variety of roles from providing nitrogen through fixation^2^ to causing disease^3^. Most of these microbes were identified in scleractinian corals from the photic zone, while studies investigating the roles of microbes in octocorals and deep-sea corals are rarer because of the limitations that great depths impose on sampling and experimentation. Such work requires the use of remotely operated vehicles or submersibles launched from ships. Still, studies have demonstrated that octocorals and deep-sea corals host some of the same associates as shallow-water scleractinians such as corallicolid apicomplexans^4–7^ and *Endozoicomonas*^8–11^. Deep-sea octocorals also host associates that are rare or absent in shallow-water and/or scleractinian corals. This includes bacteria from the SUP05 cluster whose role is linked to cold seeps in the deep sea^12^ and commonly members of the class Mollicutes.

Members of the class Mollicutes associate with many coral species and with a wide diversity of plant, fungal, and animal hosts including humans^13,14^. Several members are well-studied parasites^15^, while the impact of others on their hosts are unclear such as the ubiquitous intracellular symbionts of arbuscular mycorrhizal fungi^16–18^. Yet others are mutualists such as *Spiroplasma* spp. that infect insects and confer protection against nematodes, parasitoid wasps, and fungi through the production of unique toxins^19^. Reflecting their lifestyles as symbionts, their evolutionary history is dominated by genome reduction with some species considered to have the most reduced genomes capable of supporting cellular life^20,21^. Recently, novel and divergent mollicutes were discovered in diverse marine invertebrates including a deposit-feeding holothurian from a deep-sea trench^22^; pelagic, photosynthetic jellyfish^23,24^; wood-boring, deep-sea chitons^25^; ascidians from a coastal lagoon^26^, and crown-of-thorns sea stars from the Great Barrier Reef^27^. However, the impacts these symbionts have on their hosts remain unclear.

Among coral species, mollicutes are most commonly found in octocorals^28–37^, but have also been reported in black corals^38^ and stony corals^39,40^. These corals occupy a wide breadth of habitats from shallow-water reefs^34,37,41^, through the mesophotic zone^38^ and the deep sea^29,30,39^ to abyssal depths^42^. In many of these coral species, *Mycoplasma* spp. are the dominant microbes, comprising more than 50% of their associated microbial communities^30,37,41^. Interestingly, the relative abundances of *Mycoplasma* vary substantially across the ranges of individual coral species^36,41^. Thus, this association may be influenced by environmental conditions or otherwise shaped by geography such as through limited dispersal. In addition, *Mycoplasma* spp. exhibit varying degrees of host specificity^31,36,37^. In cases with high specificity, *Mycoplasma* spp. may have coevolved with their coral hosts and closely interact with them as symbionts. Despite these insights, the interactions between mollicutes and their host corals remain unknown.

Here we discovered two novel bacteria of the class Mollicutes which are abundant in the deep-sea octocoral *Callogorgia delta. C. delta* is common along the continental slope in the Gulf of Mexico between 400 m and 900 m depth, where it is often the dominant habitat-forming coral species^43,44^. Like many deep-sea coral species, *C. delta* colonies create habitat for numerous animal species including *Asteroschema* ophiuroids and the chain catshark, *Scyliorhinus retifer,* which lay their eggs directly on *C. delta* colonies^45^.

To characterize the association between *Callogorgia delta* and these newly discovered mollicutes, we screened colonies from six sites in the Gulf of Mexico across three years of sampling (Fig 1a) using 16S metabarcoding to determine the prevalence of these novel mollicutes and quantify their relative abundances. We also screened the closely related species *Callogorgia americana* to assess the phylogenetic breadth of the association*. C. americana,* is also a dominant habitat-forming species in the northern Gulf of Mexico and provides a good comparison species. *C. americana* occurs at shallower depths (300 – 400m) while *C. delta* is often found near hydrocarbon seeps^44^.

**Fig. 1:**
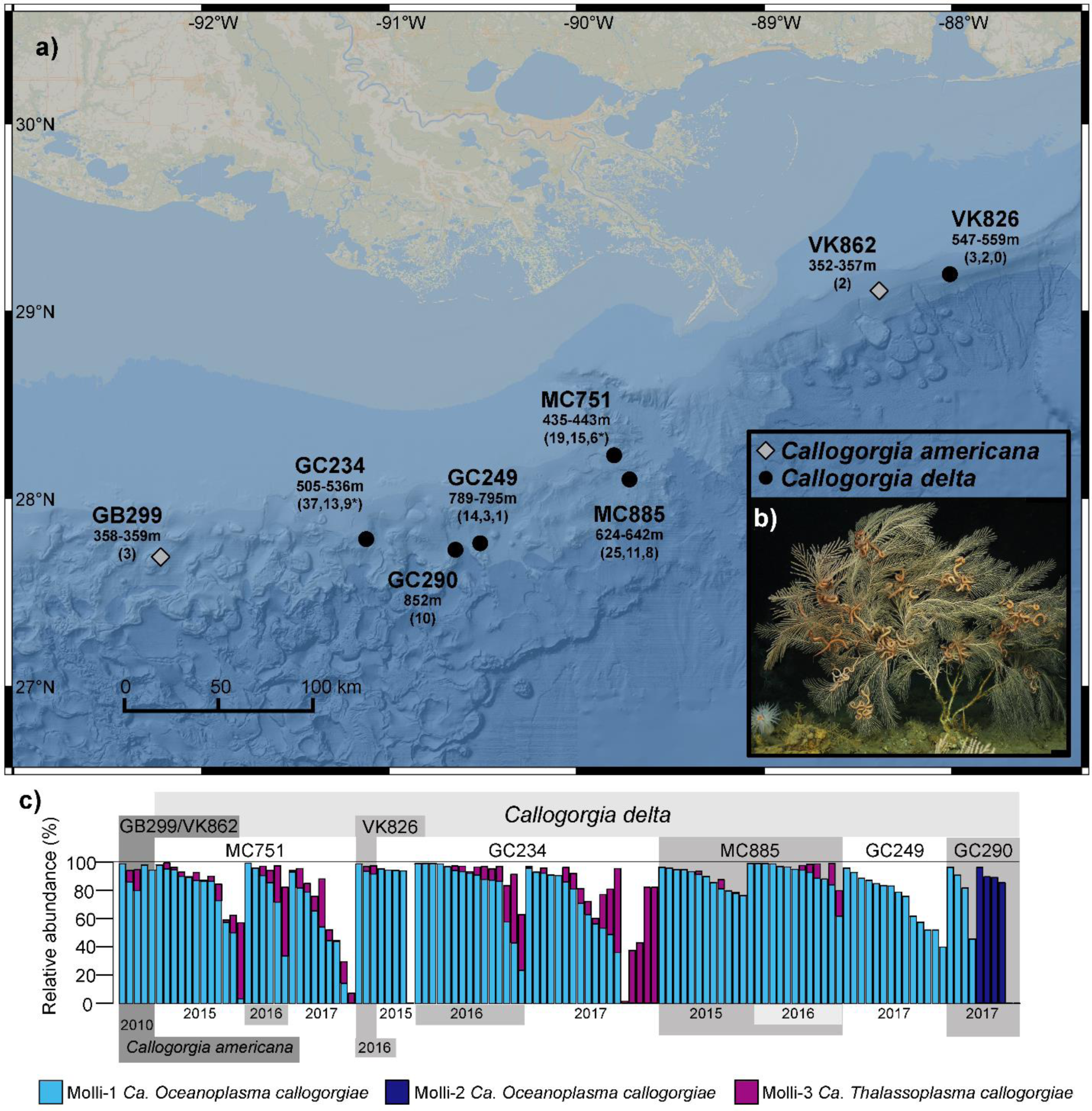
Sampling locations and prevalence of novel mollicutes in *Callogorgia delta* and *C. americana* a) Map of locations where *Callogorgia delta* (black circles) and *C. americana* (gray diamonds) were sampled. The depth range of samples collected from each site is denoted below the site name followed by the number of colonies, sediment, and water samples in parentheses. If no water or sediment samples were collected, only the number of colonies is shown. A star signifies whether a water sample was collected using a McLane pump (∼400L) in addition to niskin bottles (∼2.5L). Bathymetric map was obtained from BOEM (credit: William Shedd, Kody Kramer). b) Image of *Callogorgia delta.* c) The relative abundances of novel mollicute ASVs. Each column represents the microbial composition based on 16S rRNA libraries obtained from a single colony. Colonies are organized by species, site, and sampling year. *C. delta* sites are ordered left to right by increasing depth.

We determined the specificity of the association to corals by screening sediment and water in addition to *Callogorgia*. Further, we assembled the genomes of these novel mollicutes to assess their metabolic capabilities, determine their phylogenetic positions, and compare them to other mollicutes. Finally, we located these mollicutes within coral tissue using catalyzed reporter deposition fluorescence in situ hybridization (CARD-FISH) microscopy.

## Results

### Occurrence of novel mollicutes in and around Callogorgia

We investigated symbiont diversity using V1-V2 16S rRNA-based metabarcoding on an Illumina MiSeq. Three abundant amplicon sequence variants (ASVs) were identified among 108 *Callogorgia delta* colonies that belong to the class Mollicutes (Molli-1,2,3). Molli-1 was the most prevalent ASV (Fig. 1b). It was detected in 99 out of 108 colonies and was present in samples from all three sampling years (2015-2017) and all six sampling locations (Fig. 1b). It had an average relative abundance of 78% among corals which harbored it and reached a maximum of 99% of the microbial community in some samples. It was also present in all five *Callogorgia americana* colonies analyzed where it averaged 91% of the microbial community. Molli-2 differed from Molli-1 by a single base pair among 300. This closely related ASV was detected in 4 of 10 colonies from the deepest site, GC290. Among those four colonies, Molli-2 averaged 90% of the microbial community while Molli-1 was undetected (Fig. 1b).

Molli-3, which was 72% identical to Molli-1, was prevalent and abundant in some colonies, but less so compared to Molli-1 (Fig. 1b). Molli-3 was present in 69 out of 108 *C. delta* colonies (up to 82% of the community) and three out of five *C. americana* colonies (up to 15%). However, Molli-3 was absent in all *C. delta* colonies from the deepest sites GC249 (n=14) and GC290 (n=10).

None of these ASVs were detected in any water sample (n=30). However, Molli-1 and Molli-3 were detected in 35 and 11 out of 47 total sediment samples with maximum relative abundances of 1.1 and 0.16% of the sediment microbial community respectively.

### Phylogenetic positions of the mollicute in Callogorgia delta

Twenty-two full length 16S rRNA gene sequences that correspond to Molli-1 were assembled from separate *C. delta* metagenomes and three additional sequences were assembled that correspond to Molli-3. These sequences were distinct from those of other known mollicutes and clustered with sequences from recently discovered associates of marine invertebrates (UFBoot 100%, Fig. 2a). Interestingly, the two novel mollicutes in *Callogorgia delta* are not each other’s closest relative and shared only 84% sequence identity across the entire 16S rRNA gene which is below the cutoff which has been suggested to differentiate bacterial families^46^.

**Fig. 2:**
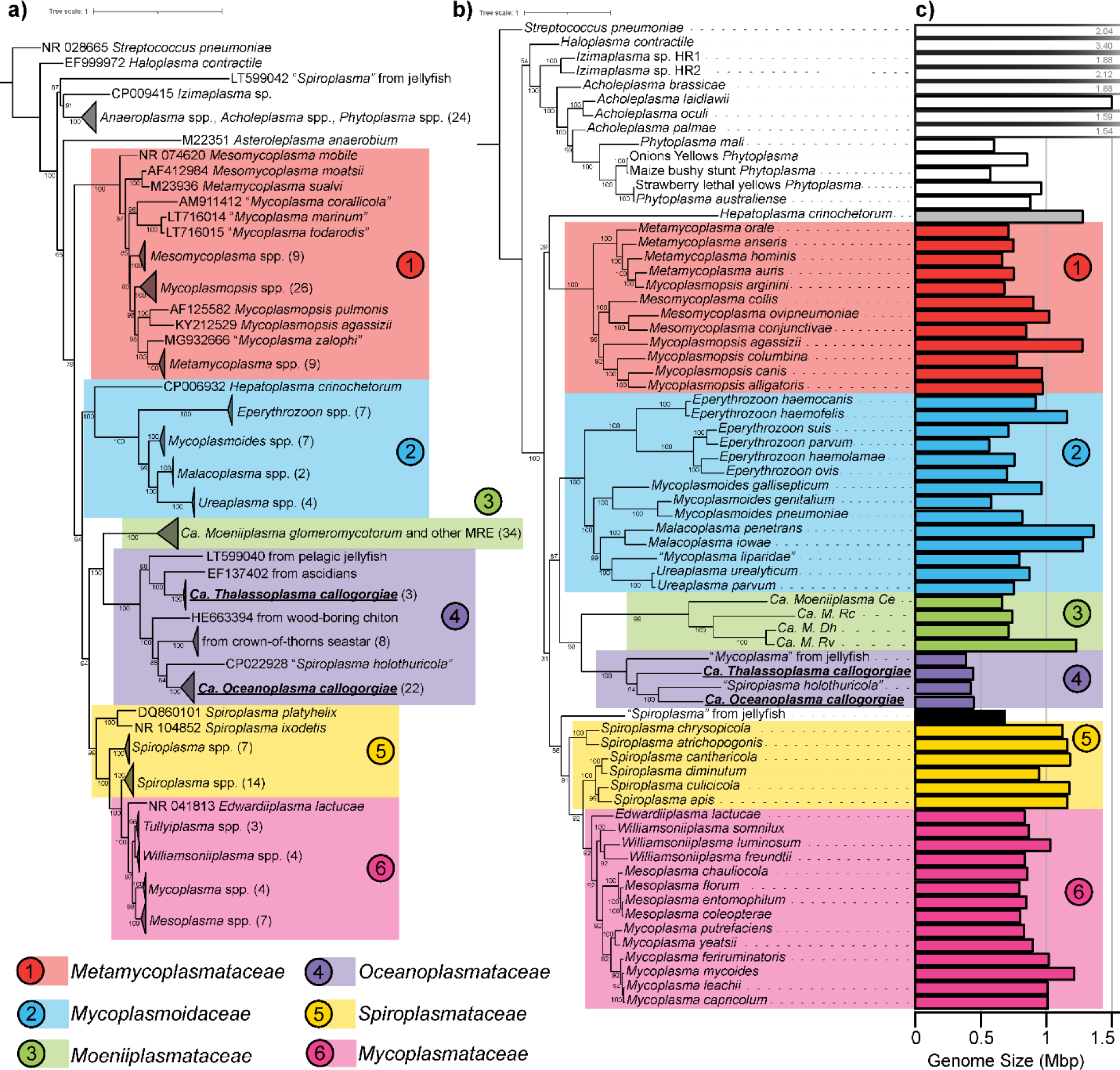
Phylogenetic analyses of the novel mollicutes in *C. delta* Maximum likelihood phylogenetic trees of class Mollicutes using the a) 16S rRNA gene and b) amino acid sequences of 43 genes. UFBoot support values are reported at each node. Numbers in parentheses designate the number of sequences within collapsed nodes. *Ca. Oceanoplasma callogorgiae* and *Ca. Thalassoplasma callogorgiae* are presented in bold text and underlined. Genome sizes (in Mbp) are depicted in c).

A circular and closed genome corresponding to Molli-1 was assembled from a deeply sequenced metagenome of *Callogorgia delta.* The metagenome was filtered to kmers with a depth of 1,000 or greater and the corresponding reads were subsampled at a rate of 10% and assembled with SPAdes^47^ to produce a single circular scaffold. Additionally, a genome corresponding to Molli-3 was obtained by co-assembling the two metagenomes with the highest 16S coverage and binning contigs using coverages from all 26 metagenomes. Phylogenomic analysis of shared amino acid sequences among mollicute genomes reiterated phylogenetic inferences based on 16S rRNA (Fig. 2b). A novel clade of Mollicutes that associates with marine invertebrates was well-supported (UFBoot 100%) and this clade clustered alongside the intracellular symbionts of mycorrhizal fungi with high confidence (UFBoot ≥98%). Branch lengths between members of the novel clade of marine mollicutes were long and the average nucleotide identities (ANI) between the four representative genomes ranged from 63 to 65% (OrthoANI^48^) suggesting that both mollicutes described here are novel genera. We propose the names *Candidatus Oceanoplasma callogorgiae* gen. nov. sp. nov. and *Ca. Thalassoplasma callogorgiae* gen. nov. sp. nov. corresponding to Molli-1 and Molli-3 respectively. Further, we erect the novel family Oceanoplasmataceae with *Ca. Oceanoplasma callogorgieae* as its type species and which includes other associates of marine invertebrates.

### Metabolic capabilities of the novel mollicutes in Callogorgia delta

The genome of *Ca. Oceanoplasma callogorgiae* was highly reduced in size (446,099 bp, Table 1, Fig. 2c) and function with only 359 predicted protein-encoding genes (241 with KEGG orthology annotations). The genome lacked any pathway to generate biomass or energy from carbohydrates and instead contained the genes comprising the arginine dihydrolase pathway (*arcABCD*) and ATP synthase. These genes were among the most highly expressed genes in both RNA libraries (Fig. 3ab) and constitute the sole pathway in this genome that is capable of producing ATP. This pathway generates ATP from the equimolar catabolism of arginine. In other non-fermentative mollicutes, additional ATP is generated as the resulting ammonia establishes a proton gradient which drives ATP synthase^49,50^. Besides arginine catabolism, a few genes could potentially be involved in ATP production: glyceraldehyde-3-phosphate dehydrogenase (*gapA*), phosphoglucomutase (*pgm*), ribose-phosphate pyrophosphokinase (*prsA*), and acetate kinase (*ackA*).

**Table 1.** Summary of the genome statistics for the novel mollicutes in *Callogorgia delta* and their closest known relatives. *LSU rRNA fragments. Pegs (w/KO) = CDS = coding sequences

| Organism name | Genome Size (bp) | Contigs | GC (%) | DRAM | Classic RAST |  |  | checkm |  |  |  |
| --- | --- | --- | --- | --- | --- | --- | --- | --- | --- | --- | --- |
|  |  |  |  | pegs (w/ KO) | CDS | tRNAs | rRNAs | Completeness (%) | Contamination (%) | Predicted Genes | Coding Density (%) |
| *Ca. Oceanoplasma callogorgiae* | 446,099 | 3 | 31.0 | 359 (241) | 405 | 34 | 2 + 7* | 70.4 | 2.6 | 392 | 88.2 |
| *Ca. Thalassoplasma callogorgiae* | 445,436 | 21 | 24.9 | 385 (237) | 416 | 30 | 2 | 72.4 | 2.6 | 396 | 87.6 |
| *Ca. Spiroplasma holothuricola*<sup>1</sup> | 424,539<sup>1</sup> | 2<sup>1</sup> | 29.5<sup>1</sup> | 336 (224) | 399 (347) | 32 | 2 | 67.4 | 2.3 | 367 | 89.9 |
| *Mycoplasma*-like from jellyfish<sup>2</sup> | 389,210 (409,158)<sup>2</sup> | 6 (11)<sup>2</sup> | 33.1 (32.9)<sup>2</sup> | 342 (225) | 358 | 32 | 2 | 73.4 (74.1) | 2.3 (2.3) | 356 | 92.5 |

References: <sup>1</sup>He et al. 2018, <sup>2</sup>Viver et al 2017

The genome featured a very limited ability to synthesize essential compounds. It encoded only one enzyme involved in amino acid synthesis (threonine dehydrogenase) but encoded several protein-degrading enzymes and amino acid importers. Similarly, it encoded no enzymes that synthesize vitamins, cofactors, or coenzymes besides NAD^+^ as well as FMN and FAD from imported riboflavin (Fig. 3a).

**Fig. 3:**
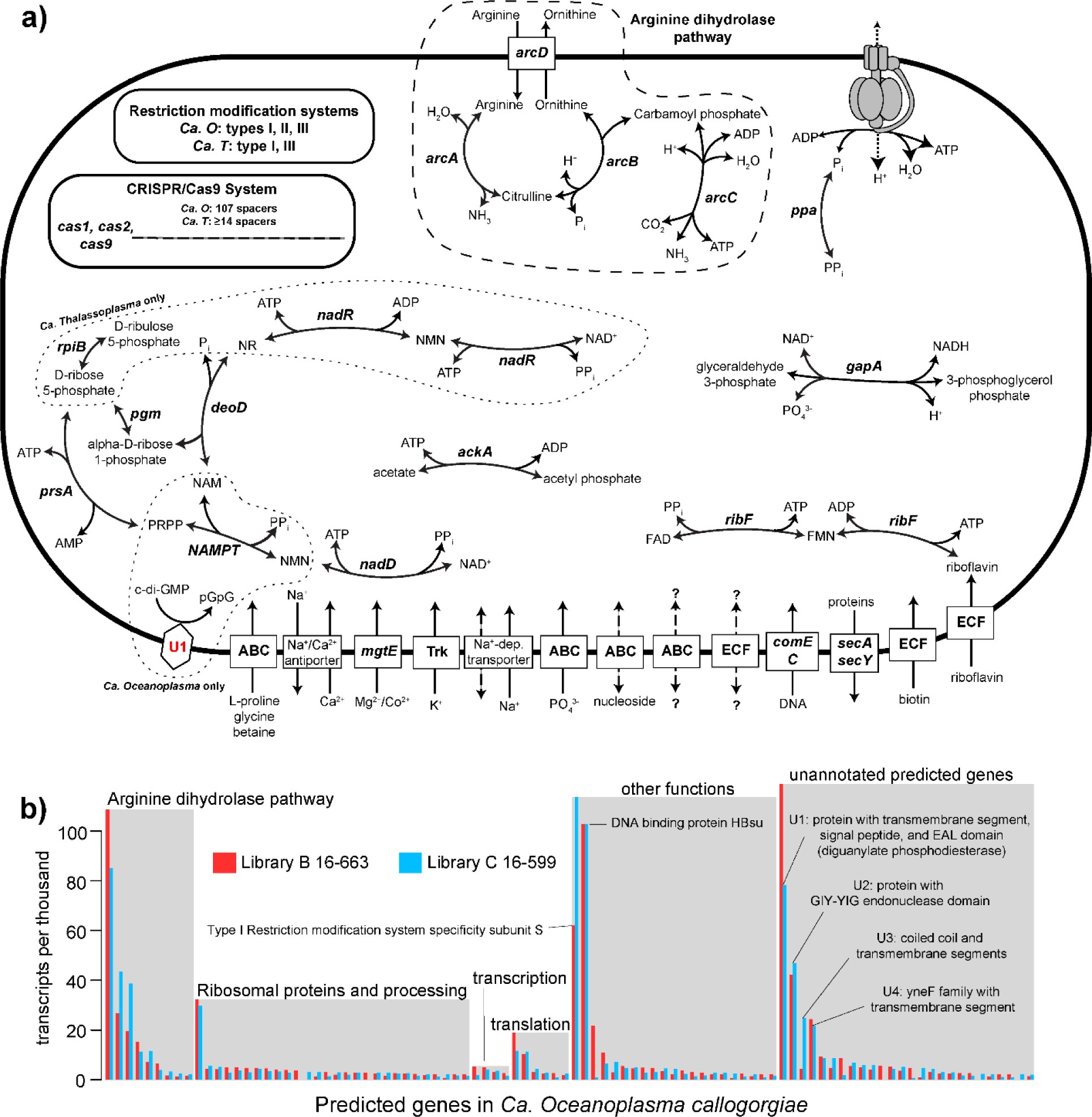
Simplified metabolic model and transcriptomic profile of novel mollicutes a) Simplified metabolic model of *Ca. Oceanoplasma callogorgiae* and *Ca. Thalassoplasma callogorgiae.* Gene names are denoted in italics and unidentified proteins in red text. ATP is generated by the arginine dihydrolase pathway. Additional ATP is likely generated by ATP-synthase utilizing the proton gradient resulting from the consumption of H^+^ and production of NH_3_. Nicotinamide (NAM), nicotinamide riboside (NR), nicotinamide mononucleotide (NMN), 5-phosphoribosyl diphosphate (PRPP) b) Expression level of *Ca. O. callogorgiae* genes from two libraries sorted by functional annotations. Only genes with a maximum of 1.5 transcripts per thousand among both libraries are displayed.

The genome contained extensive and active mechanisms to interact with foreign DNA including type I, II, and III restriction modification systems and a Class 2 Type II-C CRISPR-Cas system. One of the five most highly expressed genes in both libraries was the specificity subunit S of its type I restriction modification system (Fig. 3b). Further, it contained the second highest number of spacer sequences, at 107, of all mollicutes with available genomes^51,52^.

Four predicted genes with no functional annotations were among the ten most highly expressed (Fig. 3b). The highest of these (named U1) was predicted to contain a transmembrane segment and a signal peptide (Hhblits^53^). It also contained a region that matches the EAL domain of various proteins. The EAL domain is a diguanylate phosphodiesterase which degrades the second messenger cyclic di-GMP^54–56^. Another highly expressed, unannotated gene (U2) was predicted to contain a GIY-YIG endonuclease domain with high confidence (>95%, Hhblits). The fourth most highly expressed, unannotated gene (U4) belongs to the YneF family whose function is unknown but is conserved across Mollicutes and most Bacilli where it is highly expressed and considered an essential protein^57^.

The genome of *Ca. Thalassoplasma callogorgiae* was very similar to that of *Ca. O. callogorgiae.* It was reduced in size (445,456 bp) and metabolic capabilities with only 385 predicted protein-coding genes (237 with KEGG orthology annotations). It possessed all the pathways and genes mentioned above for *Ca. O. callogorgiae* except it lacked a type II restriction modification system and homologs of U1 and U2. In contrast, it possessed ribose-5-phosphate isomerase (*rpiB*) and alternative genes to synthesize NAD^+^ (Fig. 3a).

### Comparison to other mollicutes

The genomes of *Ca. O. callogorgiae* and *Ca. T. callogorgiae* were similar to others within Oceanoplasmataceae. The genome of “*Ca. Spiroplasma holothuricola”* is publicly available and we asssembled a fourth genome corresponding to the closely related associate of the jellyfish *Cotylorhiza tuberculata*^23^. We assembled the publicly available metagenome with the highest 16S coverage and used all metagenomes from the study for binning. These four genomes shared 211 clusters of orthologous genes (COGs) which comprised 60-66% of the COGs in each genome. Besides housekeeping genes, all four genomes encode the arginine dihydrolase pathway and CRISPR-Cas systems. The genome of the jellyfish associate was the most divergent and may have further but limited means to generate ATP since it encoded lactate dehydrogenase, glycerate kinase, ribulose-phosphate 3-epimerase, and an agmatine/putrescine antiporter.

Overall, the genomes of Oceanoplasmataceae were distinct from other mollicutes and were the most reduced (Fig. 4). They formed a distinct cluster based on the presence/absence of COGs (Fig. 4a), were the smallest (389 – 446 kbp versus >564 kbp^58^; Fig. 3c), and had the fewest protein-encoding genes (pegs) (336 – 385 versus >515; Fig. 4b). Further, they were the only mollicutes missing the genes necessary to conduct glycolysis besides *Moeniiplasma* spp. with which they shared other physiological similarities. Compared to other mollicutes, they had the fewest COGs with KEGG orthology terms and the fewest genes involved in carbohydrate metabolism. Oceanoplasmataceae genomes were distinguishable from those of *Moeniiplasma* spp. by the presence of genes encoding ribose-phosphate pyrophosphokinase, glyceraldehyde 3-phosphate dehydrogenase, and ATP synthase.

**Fig. 4:**
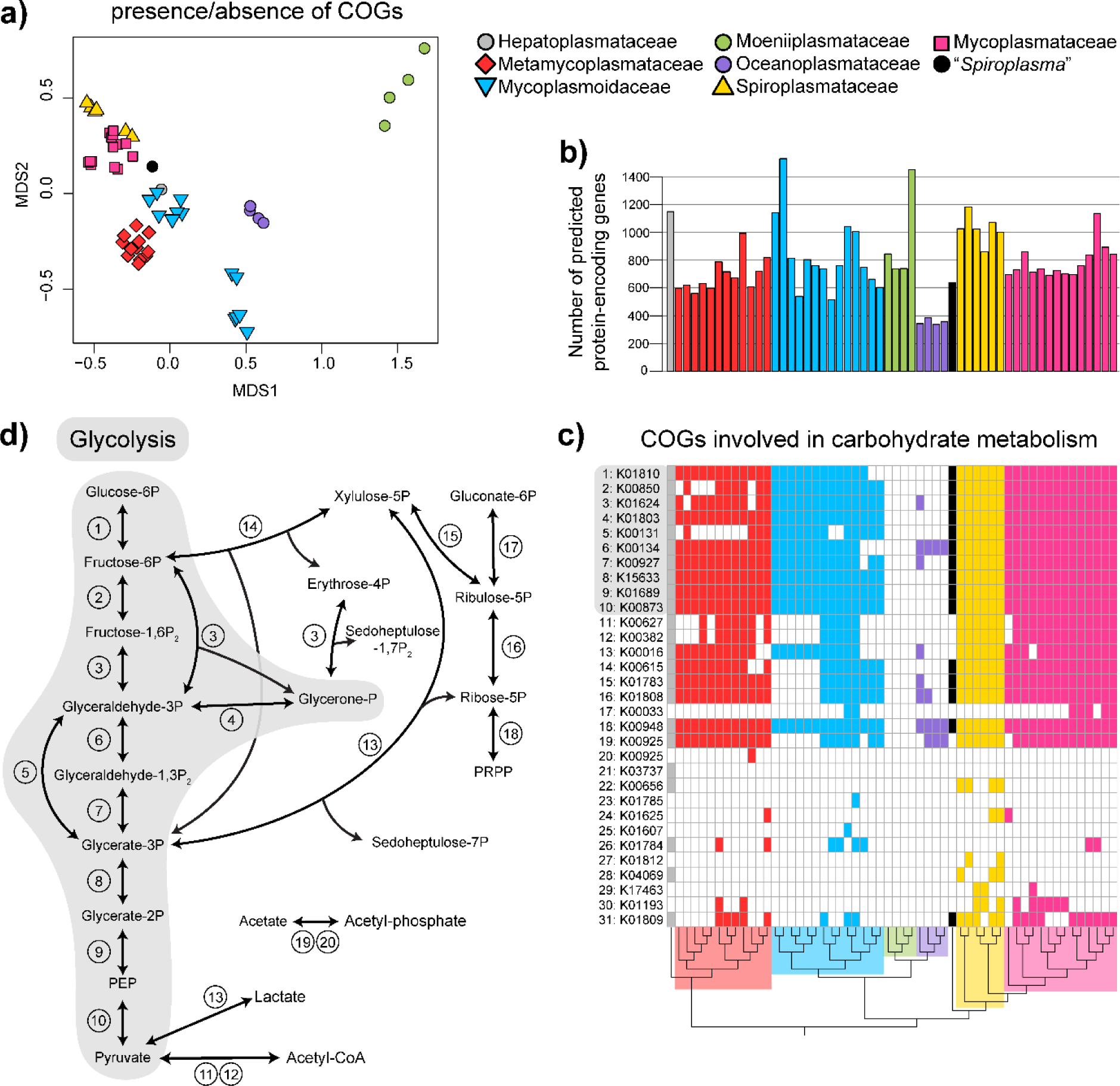
Comparative Genomics of related mollicutes a) Non-metric multidimensional scaling (NMDS) plot of cluster of orthologous genes (COG) using Jaccard distances. Only COGs present in more than two genomes were included. b) the number of predicted protein-encoding genes for each genome. Oceanoplasmataceae had the fewest predicted protein-encoding genes. c) the presence of COGs involved in carbohydrate metabolism for each genome. COGs are labeled using their KEGG orthology identifier annotations. d) metabolic map of glycolysis and other reactions associated with the COGs in c). The map was drawn using reactions listed on the KEGG database for each KO identifier. Phosphoribosyl diphosphate

All Oceanoplasmataceae genomes contained twenty-three COGs that were absent in all other mollicutes. Notably, these included a Ca^2+^/Na^+^ antiporter, several other transporters, and a thymidine kinase. Thymidine kinases were enriched in the genomes of Oceanoplasmataceae which each contained four copies while all other genomes had one or zero except two *Moeniiplasma* spp. which had two copies.

### Localization of bacteria in Callogorgia delta

To locate the symbiotic bacteria, we used probes that bind to ribosomal RNA permitting deposition of a fluorophore through catalyzed reporter deposition fluorescence in situ hybridization (CARD-FISH). Fluorescence was observed in the mesoglea of two separate *Callogorgia delta* colonies representing abundant bacteria (Fig. 5 and 6). These signals were present in tissue sections hybridized with the EUB338 I-III probe mix but absent in adjacent sections hybridized with the negative control probe (non-EUB). These signals were <1 µm in diameter and overlapped DAPI fluorescence. These signals of bacterial symbionts were observed in both the mesoglea within polyps (Fig. 5c,e,g) and in mesoglea within the coenosarc surrounding the proteinaceous central axis (Fig. 5a). They formed large aggregates potentially comprised of hundreds or thousands of cells (Fig. 6). Another fluorescence signal was observed in the base of the tentacles in one colony but was far less aggregated (Fig. 6c,d).

**Fig. 5:**
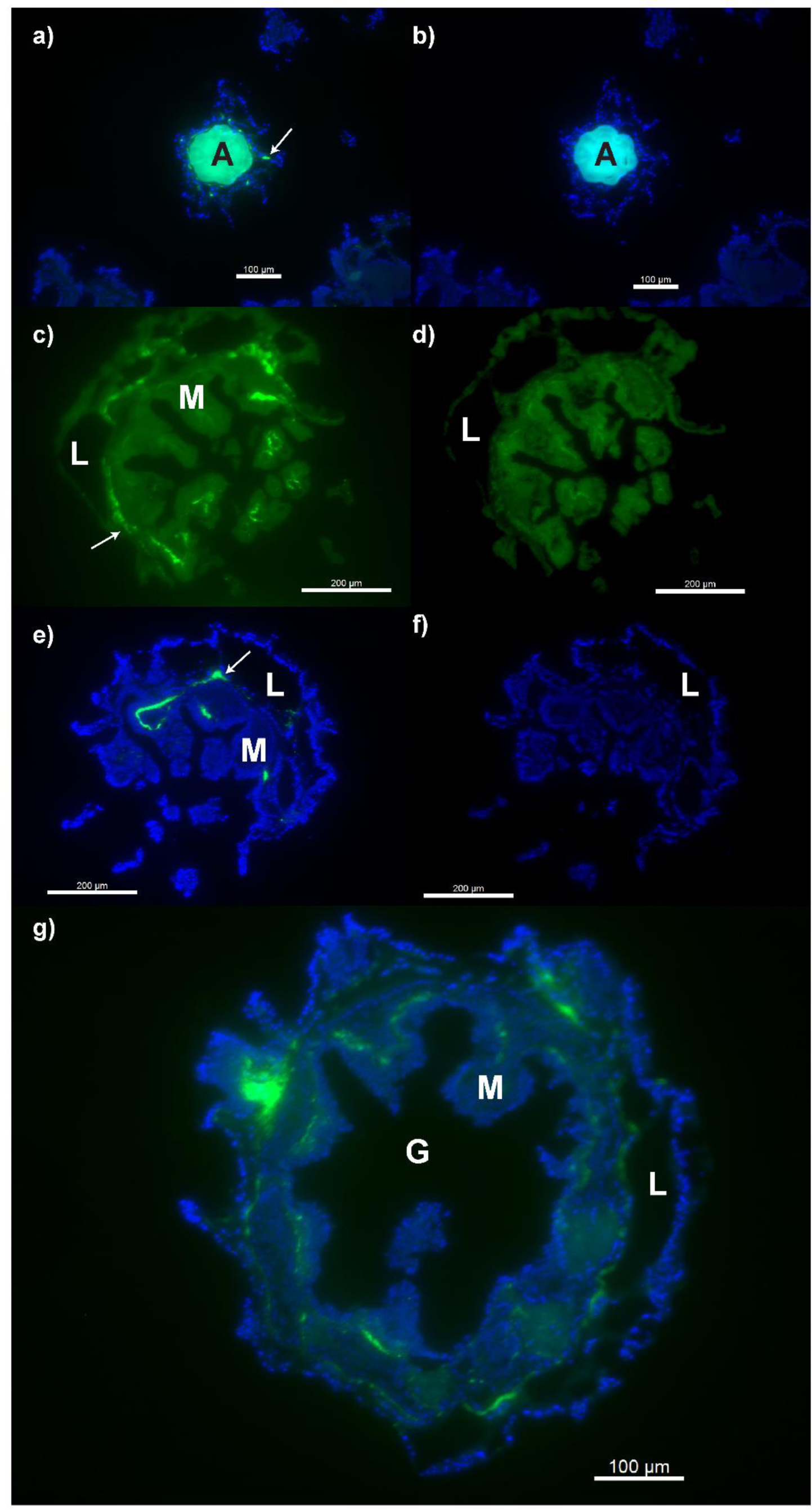
Bacterial signals in the mesoglea of *Callogorgia delta* Tissue sections were hybridized using CARD-FISH with the general eubacterial probe mix (EUB338 I-III) and DNA was stained with DAPI. Fluorescence in the emission range of DAPI is shown in blue while green shows emission in the range of the fluorophore ALEXA 488 used by both EUB338 I-III and control probes. Paired images compare adjacent sections where sections on the left were hybridized with EUB338 I-III and show bacterial signals (a,c,e) while sections on the right were hybridized with a negative control probe and only show autofluorescence (b,d,f). Signals were observed in transverse sections through the central axis (a,b) and transverse sections through a single polyp (c,d,e,f,g). Arrows in (a) and (c) show examples of EUB338 I-III fluorescence. A = central axis, G = gastrovascular cavity, M = gastric mesentery, L = sclerite lacuna. Scale bars = 100 µm (a,b,g), 200 µm (c,d,e,f)

**Fig. 6:**
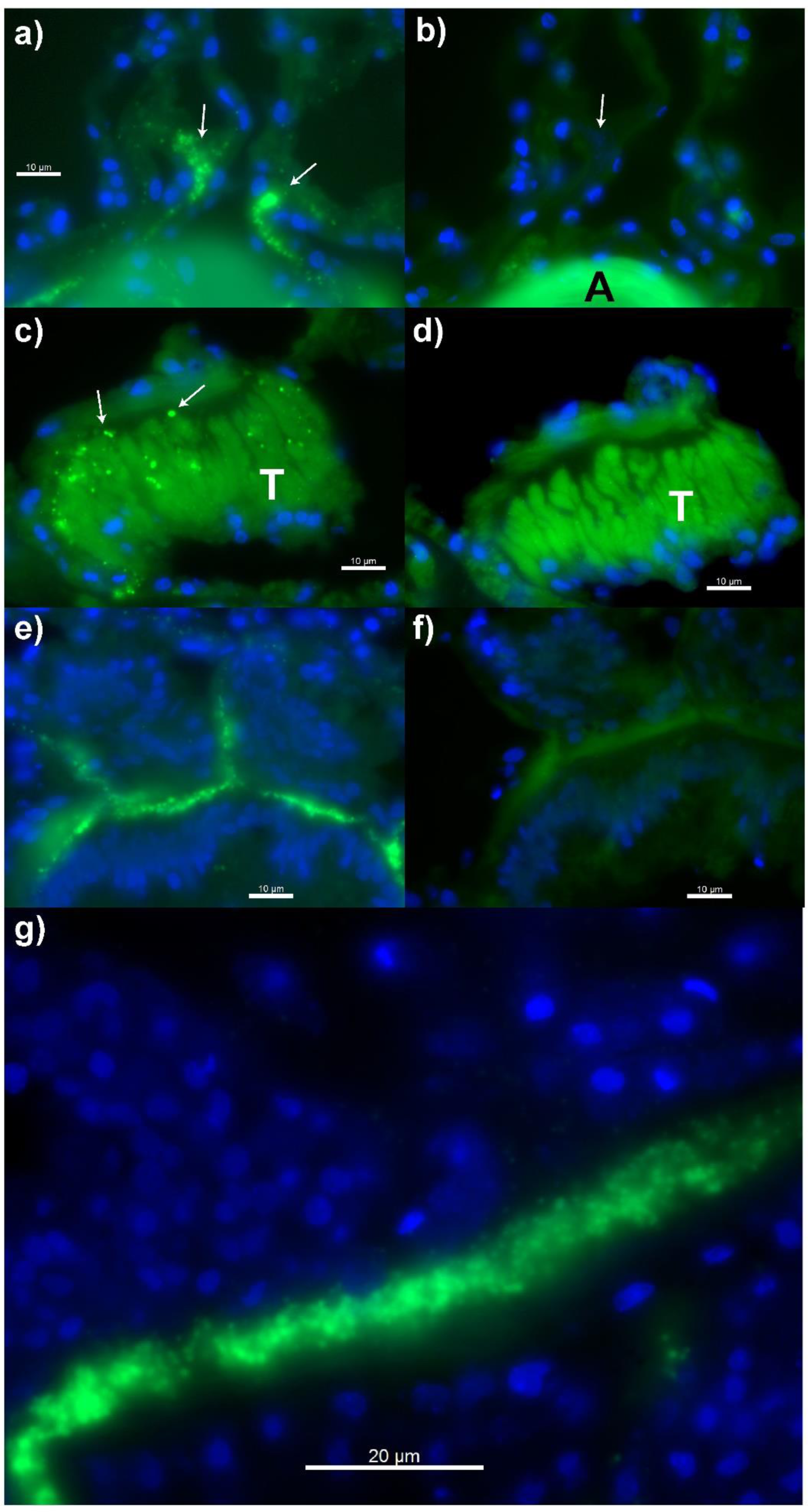
Higher magnification of bacterial signals in the mesoglea and tentacles of *Callogorgia delta* Tissue sections were hybridized using CARD-FISH with the general eubacterial probe mix (EUB338 I-III) and DNA was stained with DAPI. Fluorescence in the emission range of DAPI is shown in blue while green shows emission in the range of the fluorophore ALEXA 488 used by both EUB338 I-III and control probes. Paired images compare adjacent sections where sections on the left were hybridized with EUB338 I-III and show bacterial signals (a,c,e) while sections on the right were hybridized with a negative control probe and only show autofluorescence (b,d,f). Transverse sections through the central axis (a,b) exhibit EUB338 I-III fluorescence (a, arrows) in the surrounding mesoglea and an overlap of DAPI (c, arrow). (a, arrow). Transverse sections through a polyp (c,d) show EUB338 I-III fluorescence (c, arrows) in the base of a tentacle. Transverse sections through the body wall of a polyp show granular (<1µm) EUB338 I-III fluorescence in the mesoglea (e,f,g). A = central axis, T = tentacle. Scale bars = 10 µm (a,c,d,e,f), 20 µm (g)

## Discussion

### Novel mollicutes in Callogorgia delta are symbionts

The data presented here suggests that *Ca. Oceanoplasma callogorgiae* and *Ca. Thallasoplasma callogorgiae* are symbionts of *Callogorgia delta sensu*^59^. First, their association was stable across geography and time. Additionally, these microbes seem to associate specifically with corals in the genus *Callogorgia* since both were absent or rare in the water and sediment but were also detected in the closely related species *C. americana*. Further, their genomes were very reduced and demonstrated a reliance on compounds like arginine and amino acids that are probably obtained from *C. delta.* Finally, *Ca. Oceanoplasma callogorgiae* likely resides within the coral. Abundant bacteria were observed in the mesoglea of *C. delta* and *Ca. O. callogorgiae* was the most abundant bacterium by far in both the amplicon dataset and the metagenomes where the genomic coverage of the symbiont was up to 50 times as high as that of the host.

### Metabolic processes and other fundamental activities of the symbionts

Both *Ca. Oceanoplasma callogorgiae* and *Ca. Thalassoplasma callogorgiae* lacked genes encoding complete fermentative pathways and depended on arginine to generate ATP. They likely receive this arginine from their coral hosts as well as essential compounds such as amino acids, riboflavin, and biotin. Conversely, they export ornithine and possibly short peptides to their coral host.

*Ca. Oceanoplasma callogorgiae* possessed extensive mechanisms to defend against viruses or other forms of foreign DNA. Its genome contained an extensive CRISPR array and high expression of restriction modification systems and endonucleases. Further, it shared additional genomic features with *Ca. Thalassoplasma callogorgiae* including *comEC* to import foreign DNA and enrichment of thymidine kinase genes to salvage thymidine from DNA degradation. The presence of these anti-viral characteristics in such highly reduced bacterial genomes suggests that they play a central role in the lifestyles of these bacteria.

Other, not yet well-understood processes must underly the basic functioning of these novel mollicutes since some of the most highly expressed genes had unknown functions. This includes U1 which is likely membrane-bound and potentially degrades cyclic di-GMP. Cyclic di-GMP is a second messenger involved in morphogenesis, motility, and virulence^60^. U1 may respond to external, or host-derived, stimuli as part of signaling pathways that could be linked to host innate immunity and symbiont recognition.

### Residence in the Mesoglea

*Ca. O. callogorgiae* likely resides in the mesoglea of *C. delta.* The mesoglea of cnidarians is composed of a largely acellular, mucoid matrix of collagen containing some cells such as amoebocytes^61^. The mesoglea provides rigidity upon which the muscles of the polyp can act but also facilitates the transport of nutrients^62,63^. Some compounds diffuse through the mesoglea like glucose and fatty acids^62,63^ while amoebocytes can also shuttle compounds such as amino acids^62^. *Ca. O. callogorgiae* probably acquires compounds including arginine, other amino acids, and the precursors of coenzymes such as riboflavin from the mesoglea because it lacks the genes necessary to synthesize these.

The mesoglea in cnidarians is also a site of immune defense^64^. Bacteria that reside in the mesoglea must evade the coral’s immune defenses such as phagocytic amoebocytes^64^. A few bacteria have been localized within the mesoglea of cnidarians including *Pseudomonas* and spirochetes in *Hydra* spp.^65,66^ and pathogenic cyanobacteria in the coral *Orbicella annularis* suffering from black band disease^67^. In contrast, the only mollicute previously localized in corals was *Mycoplasma corallicola* which was found on the surface of the tentacle ectoderm of *Lophelia pertusa*^39^. Unlike *M. corallicola, Ca. O. callogorgiae* likely resides within its host coral and therefore must somehow evade phagocytic amoebocytes in the mesoglea and potentially other immune defenses.

### Consequences for the coral host

It is still unclear if *Ca. Oceanoplasma callogorgiae* incurs a cost or provides any benefit to its coral host. These novel mollicutes may simply be commensals or parasites with a minimal impact on their host corals since they did not display any obvious pathology and nearly every colony was infected. Conversely, the metabolism of arginine may provide the coral with an alternative pathway to process nitrogenous waste or could function to recycle nitrogen which would be useful for deep-sea corals because they rely on a nitrogen-poor diet of marine snow^68^. The symbiont might provide protection from viruses using its CRISPR-Cas and restriction modification systems similar to what has been proposed for the symbiont of hadal sea cucumbers which was the closest relative to *Ca. O. callogorgiae*^22^. It is also possible that some of the unannotated genes are peptides that confer protection from pathogens or parasites like those that *Spiroplasma* spp. produce in insects^19^.

### The novel family Oceanoplasmataceae

We erect the family Oceanoplasmataceae which includes *Ca. Oceanoplasma callogorgiae*, *Ca. Thalassoplasma callogorgiae*, and other associates of marine invertebrates. These hosts are phylogenetically diverse comprising four animal phyla, exhibit diverse lifestyles from being photosynthetic to wood-borers, and originate from a wide range of habitats from the epipelagic zone to hadal trenches^22–26^. In addition to this, the large phylogenetic distances between members of the Oceanoplasmataceae suggests that a substantial amount of diversity remains undiscovered. Interestingly, *Ca. Moeniiplasma* spp. appear to be the closest relatives of Oceanoplasmataceae. They clustered together in the phylogenetic analyses and shared genomic similarities such as a lack of glycolysis.

Compared to other mollicutes, Oceanoplasmataceae exhibit extensive viral defense systems. All were enriched in thymidine kinase genes and the two genomes with unfragmented CRISPR arrays had more spacers than other mollicutes. Only the associate of a hadal snailfish, “*Mycoplasma*” *liparidae,* had more spacers^52^. This may reflect a general evolutionary trend among animal-microbial symbioses in marine environments where exposure to viruses is high. Similarly, the symbionts of marine sponges are enriched in CRISPR-Cas systems compared to surrounding seawater ^69,70^.

Oceanoplasmataceae possess the most reduced genomes among all mollicutes. They possess the smallest genomes which contain the fewest protein-encoding genes and lack any complete fermentative pathway. Extensive genome reduction dominates the evolutionary history of Mollicutes. Mollicute genomes are used to identify the minimum set of genes essential for cellular life^21^ and are utilized as precursors in efforts to design minimal genomes^71,72^. These synthetic genomes retain glycolysis. However, the lack of glycolysis in Oceanoplasmataceae and *Moeniiplasma* spp. suggests that these synthetic genomes could be reduced further. Our comparative genomics analysis also revealed that *Moeniiplasma* spp. lack ATP-synthase suggesting alternative gene sets may permit further reduction.

Strangely, no mollicutes except *Acholeplasma laidlawii* are known to use cyclic di-GMP and all mollicutes instead use cyclic di-AMP^73^. The presence of an EAL domain in gene U1 of *Ca. O. callogorgiae* implies that the use of cyclic di-GMP was not lost in all other mollicutes or was regained through horizontal gene transfer in some lineages.

### Conclusion

Here we describe two novel members of the Mollicutes that associate with the deep-sea coral, *Callogorgia delta,* and propose the names *Ca. Oceanoplasma callogorgiae* and *Ca. Thalassoplasma callogorgiae*. We characterize their association with *C. delta,* generate genomic resources, and produce phylogenetic and metabolic inferences advancing our understanding of coral-associated mollicutes and symbiosis in the deep sea. Further, we erect a new family, Oceanoplasmataceae, which has not yet been recognized and whose diversity remains largely uncharacterized. We show that its genome reduction exceeds what was known among Mollicutes informing their evolutionary history, the consequences of symbiosis, and the concept of minimal bacteria.

## Methods

### Collections

Collections are a subset of those described in Vohsen et al. 2020. One hundred and eight *Callogorgia delta* colonies were collected from six sites spanning over 350 km in the northern Gulf of Mexico in 2015, 2016, and 2017 (Fig. 1). The sites are named after the Bureau of Ocean Energy Management’s designations for lease blocks in which corals were sampled and include Mississippi Canyon (MC) 751 (n=19 colonies, 435-443 m, 28.193 N −89.800 W), Viosca Knoll (VK) 826 (n=3 colonies, 547-559 m, 29.159 N −88.010 W), Green Canyon (GC) 234 (n=37 colonies, 505-536 m, 27.746 N −91.122 W), MC885 (n=25 colonies, 624-642 m, 28.064 N - 89.718 W), GC249 (n=14 colonies, 789-795 m, 27.724 N −90.514 W), and GC290 (n=10 colonies, 852 m, 27.689 N −90.646 W). To compare *Callogorgia delta* to a closely related coral species, we additionally processed five *Callogorgia americana* colonies that were collected in 2010 from Garden Banks (GB) 299 (n=3 colonies, 358-359m, 27.689 N −92.218 W) and VK862 (n=2 colonies, 352-357m, 29.109 N −88.387 W).

Colonies were sampled using specially designed coral cutters mounted on the manipulator arm of remotely operated vehicles (ROVs). Coral fragments were removed, placed in separate temperature insulated containers until recovery of the ROV, and were maintained at 4 °C for up to 4 hours until preservation in ethanol or freezing in liquid nitrogen.

Sediment samples were taken in close proximity to many of the *Callogorgia delta* collections using push cores with a diameter of 6.3 cm. Upon recovery of the ROV, 1 mL of sediment from the top 1 cm of each sediment core was frozen in liquid nitrogen for later DNA extraction for microbiome analysis. Water was sampled in 2015 using a Large Volume Water Transfer System (McLane Laboratories Inc., Falmouth, MA) which filtered approximately 400 L of water through a 0.22 micron porosity filter with a diameter of 142 mm. One quarter of each filter was preserved in ethanol and another quarter was frozen in liquid nitrogen for these analyses. In 2016 and 2017, 2.5 L Niskin bottles mounted on the ROV captured water above corals. Upon recovery of the ROV, this water was filtered through 0.22 micron filters which were frozen in liquid nitrogen.

### 16S rRNA metabarcoding

The microbiomes of *Callogorgia delta, Callogorgia americana*, water, and sediment were analyzed using 16S metabarcoding. DNA was extracted from coral tissue and sediment samples using DNeasy PowerSoil kits (Qiagen, Hilden, Germany) following manufacturer protocols using approximately 1 cm of coral branches and about 0.25 g of sediment. DNeasy PowerSoil kits were also used for water samples collected in 2015 using 1 cm^2^ of filter. For all water samples from 2015, replicate extractions were performed on quarters of the filter preserved in both ethanol and frozen in liquid nitrogen. DNA was extracted from all other water samples using Qiagen DNeasy PowerWater kits.

The V1 and V2 regions of the 16S rRNA gene were amplified using universal bacterial primers 27F and 355R^74^ with CS1 and CS2 adapters (Illumina, San Diego, CA). PCR was conducted with the following reaction composition: 0.1 U/ µL Gotaq (Promega, Madison, WI), 1X Gotaq buffer, 0.25mM of each dNTP, 2.5 mM MgCl_2_, and 0.25 µM of each primer, and the following thermocycler conditions: 95 °C for 5 min; 30 cycles of 95 °C for 30 sec, 51 °C for 1 min, and 72 °C for 1 min; and finally 72 °C for 7 min. Libraries were prepared by the University of Illinois Chicago DNA services facility and sequenced on two separate runs on an Illumina MISEQ platform^75^. Raw sequence data are available on the NCBI database under BioProject PRJNA565265 and BioSample IDs SAMN12824733 – 4765, SAMN12824767 – 4875.

Amplicon sequence data were analyzed as described in^4^ using QIIME 2 (ver2017.11)^76^. Reads were joined using vsearch and quality filtered using q-score-joined. Deblur denoise was used to detect chimeras, construct sub-operational taxonomic units (referred to as ASVs), and trim assembled reads to 300bp.

### Metagenomes and Metatranscriptomes

Metagenomes and metatranscriptomes of *C. delta* were sequenced to assemble genomes of novel mollicutes which were used to assess their phylogenetic positions and metabolic potential. DNA was extracted from eight *Callogorgia delta* colonies as described in Vohsen et al. 2020. In brief, DNA was extracted using DNA/RNA allprep kits (Qiagen, Hilden, Germany) with β-mercaptoethanol added to the RLT+ lysis buffer following the manufacturer’s instructions. These DNA extracts were sequenced on an Illumina HISEQ2500 platform using 150 bp paired-end reads. The two libraries with the highest coverage of mollicutes were sequenced a second time. In addition to these, DNA was extracted from an additional 18 samples corresponding to 16 additional colonies. These were sequenced on an Illumina HISEQ platform with 100bp paired-end.

RNA was extracted from two *Callogorgia delta* colonies using an RNeasy extraction kit (Qiagen, Hilden, Germany). The RNA extracts were enriched for bacteria by depleting coral host rRNA using a Ribo-Zero Gold Yeast kit (Illumina, San Diego, CA, USA) following the manufacturer’s protocols. The RNA extracts were sequenced on a MISEQ platform with 150 bp single-end reads.

All DNA sequence libraries were screened for bacterial SSU rRNA using phyloFlash ver3.3^77^. The library with highest coverage of *Ca. Oceanoplasma callogorgiae* was used to assemble its genome. Reads were trimmed to a quality score of 2 from both ends using BBduk and a kmer frequency analysis was performed using BBnorm. The library was kfiltered to a depth of 1,000 or greater using BBnorm and downsized to 10% using reformat with a single pass. The library was then assembled using SPAdes^47^ with a kmer progression of 21,33,55,77,99,127. Bandage ver0.8.1^78^ was used to identify a single circular scaffold of contigs containing the 16S rRNA gene of *Ca. Oceanoplasma callogorgiae*. The overlaps on the end were manually trimmed off and the scaffold was recentered to the origin of replication predicted by Ori-Finder^79^. This genome was then annotated using classic RAST ver2.0^80,81^.

In order to generate a genome for *Ca. Thalassoplasma callogorgiae*, the two libraries with the highest coverage of its 16S rRNA gene were coassembled using megahit^82^. All libraries were mapped to this coassembly using BBmap with a kfilter of 22 bp, subfilter of 15 bp, and a maxindel of 80 bp. Alignments were converted from SAM to BAM formats, sorted using samtools^83^, and used for binning using metabat^84^ with default parameters. All scaffolds with contigs belonging to bin one constituted a draft genome that was annotated using RAST and used for phylogenomic analysis.

The presence of CRISPRs and associated genes along with the number of spacers were determined using RAST annotations and CRISPRfinder^85,86^. OrthoANI^48^ was used to estimate average nucleotide identity between genomes. To determine which genes were expressed and to quantify their expression, the two RNA sequence libraries were pseudoaligned to the DRAM gene annotations described below of both genomes with kallisto ver0.44.0^87^ using 100 bootstrap replicates, an estimated fragment length of 180 bp with a standard deviation of 20 bp, and an index with a k-mer size of 31 bp. Coding regions with high expression levels but no functional annotations from RAST were further investigated using blastp and HHBlits^53^ to inform potential function.

### Phylogenetic Analyses

Phylogenetic trees were constructed to infer the phylogenetic positions of the mollicutes that were detected in *Callogorgia delta.* First a tree of 16S sequences was generated to place these novel mollicutes within the wide diversity of mollicutes with available 16S sequences. Full length 16S rRNA genes were assembled from the metagenomes using phyloFlash ver 3.3^77^. These full-length sequences were aligned with publicly available Mollicutes sequences that were over 1,000 bp using MUSCLE ver3.8.425^88^. A maximum likelihood tree was constructed using the IQ-TREE web server ver1.6.10^89,90^. Within IQ-TREE, ModelFinder^91^ was used to choose a general time reversible model with unequal rates and empirical base frequencies as well as a FreeRate model^92,93^ for rate heterogeneity across sites with 3 categories (GTR+F+R3) based on the highest Bayesian information criterion (BIC) score. UFBoot2^94^ was used to obtain ultrafast bootstrap support values (UFBoot) using 1000 replicates.

To obtain a more confident phylogenetic placement of the coral-associated mollicutes, a phylogenomic tree was constructed including genomes from the two mollicutes in *C. delta* and 66 mollicute genomes accessed through GenBank and the European Nucleotide Archive (ENA) (Supplementary File 1g). One more mollicute genome was assembled from a dataset available on ENA because an assembled genome was not available. This genome corresponded to the novel bacterium that associates with the jellyfish, *Cotylorhiza tuberculata*, which was tentatively identified as *Mycoplasma*^23^. Among the four libraries, M3 was chosen for assembly since it had the highest coverage of *Mycoplasma* based on the coverages of 16S rRNA assembled using phyloFlash^77^. This library was then assembled using megahit^82^ with a kmax of 241. All four libraries were mapped to this assembly using bbmap with a kfilter of 22 bp, subfilter of 15 bp, and maxindel of 80 bp. The alignments were subsequently used to bin the assembly using metabat^84^ using default parameters. Bin one corresponded to the purported *Mycoplasma* genome. This bin and all associated scaffolds were used to represent this genome in phylogenomic analyses and was also annotated using RAST to compare to other genomes. A concatenated amino acid sequence alignment of 43 genes from all 69 genomes was generated with CheckM (ver1.1.3)^95^. A maximum likelihood phylogenomic tree was then constructed using IQ-TREE (ver1.6.12)^89,94^. Both phylogenetic trees were annotated using the recently revised phylogeny of the class Mollicutes^96–98^.

#### Comparative Genomics Analyses

All mollicute genomes included in phylogenetic analyses that use translation table 4 (all except Acholeplasma-Anaeroplasma-Phytoplasma group) were annotated using DRAM^99^. Clusters of orthologous genes (COGs) were identified among all genomes using orthoMCL^100^ on KBase^101^ with the “BuildPangenome with OrthoMCL” app (version 2.0) using amino acid translations from all protein-encoding genes (pegs) and default parameters. To visualize differences in genomic composition, non-metric multidimensional scaling was performed using Jaccard distances based on the presence/absence of COGs. Only COGs that were present in more than two genomes were included.

### Microscopy

CARD-FISH was employed to localize bacteria within *Callogorgia delta.* Colonies collected in 2016 were preserved for microscopy. During subsampling, 0.5 cm long branches were fixed in 2 mL of 4% formaldehyde with 1X phosphate-buffered saline (PBS) at 4 °C overnight (4-8 hours). Samples were washed 3 times and then stored in equal parts ethanol and 1X PBS for up to 21 months. Samples were decalcified in 0.45M EDTA in 1X PBS for 7 days, refreshing the buffer after 4 days, then washed in 1X PBS. Samples were post-fixed in 1X PBS with 1% paraformaldehyde for 1 hour then washed 3 times in 1X PBS and stored in 50:50 1X PBS:ethanol overnight. The next day, the samples were embedded in TissueTek (Sakura Finetek, Maumee, OH) and left at 4 °C overnight. The next day, the TissueTek was replaced and the samples were frozen. Samples were sectioned to a thickness of 4 µm using a Leica (Wetzlar, Germany) cryostat CM3050 S. Adjacent sections were placed on polysine slides and stored at 4 °C before hybridization.

Hybridization began by removing TissueTek from tissue sections via soaking slides in milliQ water for 3 minutes. Autofluorescence was then diminished by soaking slides in ethanol with 0.1% Sudan black B for 10 minutes. Sections were washed three times in 1X PBS and left in 1X PBS for a final 24 minutes. Endogenous peroxidases were inactivated by soaking sections in 0.2M HCl for 12 minutes, then 20 mM Tris-HCl for 20 minutes, lysozyme solution (0.01 g/mL) for 30 minutes at 37 °C, 20 mM Tris-HCl again for 20 minutes, and finally 3 minutes in milliQ water. Slides were then airdried and sections were circled with a PAP pen. Two probes were used: a mix of EUB338 I-III which target eubacteria^102,103^ and a negative control probe (non-EUB) consisting of the reverse complements. Hybridization mixes were created by combining 1 µL of probe (50 ng/µL) with 299 µL hybridization buffer (30% formamide with 0.102 M NaCl, 0.02 M Tris HCl, 10% Blocking Reagent, 0.01% sodium dodecyl sulfate, and 0.1 g/mL dextran sulfate). Each section was covered with approximately 10-20 µL of one of the hybridization mixes. Two adjacent sections were hybridized with the EUB338 I-III probe mix and non-EUB probes separately to produce a negative control of the same tissue region. Each slide was then placed in a humidity chamber consisting of a closed 50 mL tube with a kimwipe soaked in 2 mL of 30% formamide in milliQ water. Hybridization was accomplished by incubating slides at 46 °C for 2 hours within their humidity chambers.

After hybridization, slides were washed for 15 minutes at 48 °C in washing buffer (0.102M NaCl, 0.02M Tris HCl, 0.005M EDTA, 0.01% SDS) followed by 15 minutes in 1X PBS at room temperature. A fresh amplification mix was created by first combining 299 µL 1X PBS and 1 µL H_2_O_2_, then adding 10 µL of that solution to 1 mL of amplification buffer (1X PBS, 0.01% Blocking Reagent, 2M NaCl, 0.1 g/mL dextran sulfate) and 1 µL of labelled tyramide (1mg/mL ALEXA 488-labelled tyramide in dimethylformamide with 20 mg/mL p-iodophenylboronic acid). Slides were again placed in 50 mL humidity chambers with a kimwipe soaked in 2 mL of milliQ water. Amplification was accomplished by incubating slides at 37 °C for 30 minutes in these humidity chambers.

After amplification, slides were washed by dipping in 1X PBS followed by 10 minutes in 1X PBS, and a final dip in milliQ water. Slides were air dried and sections were stained with DAPI (1 µg/mL) for 10 minutes in the dark. Slides were washed a final time in milliQ water for 3 minutes. Once airdried, slides were mounted using Vectashield (Vector Laboratories, Burlingame, CA) and viewed on an Axioscope epifluorescence microscope (Carl Zeiss AG, Oberkochen, Germany) with filters restricted to the emission spectra of ALEXA 488 and DAPI. Images were taken with a Zeiss AxioCam camera and visualized using AxioVision software. CARD-FISH signals were considered positive if they appeared consistently within a structure in the coral, were absent in the same structures in negative control sections, and overlapped DAPI signals.

## Data availability

Raw sequence data are available on the NCBI Sequence Read Archive under BioProjects PRJNA574146 (BioSample IDs SAMN12856800 – SAMN12856807) and PRJNA565265 (SRA accession numbers SRR10174410 and SRR10174411). Assembled 16S rRNA sequences and genomes are available under accession numbers OR679038 – 62, CP125803, and JARVCM000000000. Raw microscopy images are available upon request. Data used to construct figures are provided in Supplementary File 1.

## Code availability

No custom scripts were developed for this manuscript that are central to the main findings. Code used in this manuscript followed manual guidelines with details described in the methods. Scripts are available upon request from the corresponding author.

## Acknowledgements

We would like to thank Martina Meyer, Miriam Sadowski, Silke Wetzel, and Meghann Devlin-Durante for their help with microscopy and DNA extractions. We would also like to thank the crews of all research vessels and ROV teams involved in collecting samples. This research was made possible by a grant from The Gulf of Mexico Research Initiative which was awarded to the Ecosystem Impacts of Oil and Gas Inputs to the Gulf (ECOGIG) consortium. Additional funding was provided from the Max Planck Society. The funders had no role in study design, data collection and analysis, decision to publish, or preparation of the manuscript.

## Author Information

### Authors and Affiliations

Department of Biology, The Pennsylvania State University, State College, PA, USA Samuel A. Vohsen, Charles R. Fisher, Iliana B. Baums

Department of Symbiosis, Max Planck Institute for Marine Microbiology, Bremen, Germany Harald R. Gruber-Vodicka, Nicole Dubilier

### Contributions

SAV conducted sampling, processing, and analyses. IBB and CRF acquired funding, led research cruises, and supervised work. ND contributed additional funding and supervised work. HGV advised analyses. SAV wrote the manuscript. All authors reviewed and edited the manuscript.

### Corresponding author

Correspondence to Iliana B. Baums

